# Short-term cortical stimulation combined with targeted rehabilitation and Chondroitinase ABC improves upper limb function following a cervical contusion injury

**DOI:** 10.64898/2026.10.07.757376

**Authors:** Eleni Sinopoulou, Emily R. Burnside, Katalin Bartus, Elizabeth J. Bradbury

## Abstract

Spinal cord injuries (SCI) at the cervical level can lead to permanent upper limb and hand motor impairments, thus restoring upper limb function is amongst the highest priorities for SCI patients and can have extensive effects on improving quality of life and patient independence. We previously showed that long-term cortical stimulation of the motor cortex (10 weeks) when applied in conjunction with a daily upper limb targeted rehabilitation regiment and a neuroplasticity enhancing therapy such as lentiviral Chondroitinase ABC (ChABC) can lead to an increase of muscle activity which was translated to restored skilled function after a clinically relevant contusion injury. Here, we investigate the effects of a short term cortical stimulation protocol as part of the same neurorehabilitation paradigm to asses its efficecy. Adult rats were implanted unilaterally with stimulating electrodes (contralaterally to the dominant forepaw as determined by skilled reaching) on the surface of the motor cortex and received a bilateral cervical (C5/C6) contusion injury to produce severe and long lasting forelimb motor deficits. Contused animals received acute intraspinal injections of the neuroplasticity promoting agent ChABC and 12 consecutives days of epidural electrical stimulation of the motor cortex along with daily targeted upper limb behavioural rehabilitation (Whishaw Window and Montoya staircase, 15 minutes for 5 days/week for 7 weeks) starting at 7 days post injury. Our results revealed marked improvements in the tasks specific for assessing upper limb dexterity. These were also corroborated by electrophysiological and anatomical evidence. Thus, we provide robust evidence that short term cortical epidural stimulation as part of a combinational neurorehabilitation paradigm can restore upper limb function after a cervical contusion.

**Significance Statement:** This study is the first instance where a neurorehabilitation paradigm for restoring upper limb function combines cortical epidural electrical stimulation of the motor cortex and applied in conjunction with upper limb targeted rehabilitation and the neuroplasticity enhancing agent Chondroitinase ABC (ChABC). The use of a short neuromodulation approach (12 days) and a cervical contusion injury model significantly increases the clinical relevancy of these preclinical experiments.

## INTRODUCTION

Spinal cord injury (SCI) is a debilitating condition leading to chronic impairments and dysfunctions thus significantly affecting the patient’s independence and quality of life. Cervical contusion injuries are the most common type of injury and lead to severe upper limb and hand impairments therefore, restoring upper limb and hand function is amongst the highest priorities for SCI patients (Anderson, 2004). One of the affected descending pathways, the Corticospinal tract (CST) is the spinal pathway most involved in skilled motor function, thus a potential and promising target for neurorehabilitation treatment options (Xu et al., 2019).

One of the biggest challenges for restoring function after SCI is the limited capability of the CNS to regenerate and the lack of intrinsic capacity to initiate axonal growth (Ferguson and Son, 2011; Mahar and Cavalli, 2018). However, seminal studies conducted in the early 1980s demonstrated that CNS neurons do have the capacity for regeneration given a permissive environment such as the PNS environment (David and Aguayo, 1981; Richardson et al., 1980). Later studies, used the bacterial enzyme lentiviral chondroitinase ABC (LV-ChABC) to break down chondroitin sulfate proteoglycans (CSPGs) after injury and targeted the CNS intrinsic inhibition leading to regeneration and neuroplasticity, thus creating a permissive environment (Carter 2008). Results from a plethora of studies showed promising effects of ChABC after CNS injury such as promoting axonal regeneration in the adult brain after a lesion (Moon et al., 2001) and leading to functional recovery after a SCI in rodents (Bartus et al., 2014; Bradbury et al., 2002; James et al., 2015), cats (Tester and Howland, 2008) and canines (Hu et al., 2018).

Although, there is evidence supporting some degree of spontaneous recovery after SCI (Fawcett et al., 2007; Steeves et al., 2011) most patients will experience permanent functional impairments. Currently, physical rehabilitation is the most common form of treatment used in the clinic for SCI patients (Fu et al., 2016), therefore it is essential to consider any form of treatment intervention in the context of physical rehabilitation. There is an increased amount of evidence suggesting that targeted rehabilitation combined with ChABC leads to promising results such as significant recovery of skilled function (García-Alías et al., 2011; Wang et al., 2011; Wiersma et al., 2017).

Another promising intervention to promote functional recovery following a CNS injury is the use of electrical neuromodulation (James et al., 2018). Previous studies from the Martin group showed that the use of epidural cortical stimulation independently of rehabilitation demonstrated promising results in rats after a unilateral pyramidotomy injury, who received 6 hours of epidural subthreshold cortical electrical stimulation for 10 days. Furthermore, these studies showed both significant functional improvements and increased CST sprouting in the spinal cord (Carmel et al., 2013, 2010). They also demonstrated functional improvements in skilled tasks, such as the horizontal ladder crossing, which were retained for 30 days after the end of the stimulation paradigm (Carmel et al., 2010). Therefore, it has been previously demonstrated that 10 days of cortical stimulation can lead to significant functional improvements and anatomical changes after CNS injury. A recent study from the same group also showed that transcutaneous spinal cord stimulation (tcSCS) paired with Theta Burst stimulation (TBS) stimulation administered for 10 consecutives days, on rats with a C4 midline contusion, led to an increase in CST axonal growth which also translated in partial recovery of skilled motor functions (Zareen et al., 2017).

Moreover, a recent study from our group has positively demonstrate that the combination of a long term cortical epidural stimulation protocol (10 weeks) when combined with upper limb specific rehabilitation and acute intraspinal LV-ChABC injections leads to restored skilled function in reaching specific tasks and an increase in evoke and voluntary muscle activity (Sinopoulou et al, 2022).

This study is the first instance where a short term (12 days) cortical epidural stimulation protocol, intensive and targeted upper limb rehabilitation and intraspinal injections of the neuroplasticity promoting agent LV-ChABC have been combined and used as a treatment paradigm to restore upper limb function in a rat cervical contusion model. Here, we aim to test whether our combinational neurorehabilitation paradigm will be effective and able to target several SCI pathology features by; (i) Using LV-ChABC injections to create a permissive environment at the injury site, (ii) using cortical electrical stimulation to promote and increase axonal sprouting and neuroplasticity and (iii) using daily upper limb targeted rehabilitation to increase specificity and guide newly formed activity dependent connections.

## MATERIALS AND METHODS

### Animals and Experimental design

Adult female Lister Hooded rats (200 –250 g; Charles River) were used for all experiments and were single housed under a 12 h light/dark cycle with ad libitum access to food and water. All procedures were performed in accordance with the United Kingdom Animals (Surgical Procedures) Act 1986. The study consisted of the following groups: uninjured/not implanted control group (n=4; only used for terminal electrophysiology experiments), no treatment group (n=6), intraspinal LV-ChABC injections only group (n=6) and epidural stimulation plus daily behavioural rehabilitation plus intraspinal LV-ChABC injections (n=6) referred to as full treatment group. Animals were randomly assigned in the different treatment groups.

### Surgical Procedures

#### Primary motor cortex (M1) epidural electrode implantation

An epidural bipolar tungsten stimulating electrode with a length of 2 mm was used for stimulating the motor cortex (M1) over the caudal forelimb area (CFA) (MSS333/1-A/SPC; PlasticsOne, Inc). The tips of the electrode were de-insulated for better dura contact using a drummel. The rats were anaesthetized (ketamine, 60 mg/kg, and medetomidine, 0.25 mg/kg) and the back of the neck and both forelimbs were shaved and cleaned with iodine. Core temperature was maintained close to 37 °C using a self-regulating heated blanket for the duration of the procedure. The electrode was sterilized with 10% chlorox and then rinsed with 70% isopropyl alcohol and sterile saline. The head was fixed in a stereotaxic frame and a small craniotomy was drilled at the level of bregma, extending from 1 mm to 3 mm lateral to midline, and from 1mm to 3 mm rostral to midline. After the craniotomy on the M1 site, the electrode was placed on the dura over the caudal forelimb area contralaterally to the dominant forepaw, as prior determined by single pellet reaching. The electrode/craniotomy site was covered with gel foam to avoid dental acrylic contact with the dura. Finally, the electrode was stabilized with dental acrylic bridged to skull screws.

#### Contusion spinal cord injury

1 week after electrode implantation all animals underwent cervical contusion injuries, as previously described (Burnside et al., 2018; James et al., 2015). Rats were anaesthetized (ketamine, 60 mg/kg, and medetomidine, 0.25 mg/kg), the cervical region of their backs was shaved and cleaned with iodine and core temperature was maintained close to 37 °C using a self-regulating heated blanket. A laminectomy was performed at vertebral levels C5 and C6 to expose the underlying spinal cord. Animals were then positioned and stabilized in an Infinite Horizon impactor (Precision Systems Instrumentation, Lexington, KY) with the impactor probe positioned 3–4 mm above the exposed spinal cord. The impactor probe delivered an impact force of 225 kD to the spinal cord, inducing a moderate midline cervical contusion injury. Following the injury, rats designated to a treatment group that didn’t include LV-ChABC intraspinal injections had their wounds sutured and anaesthesia was then reversed using atipamezole hydrochloride (1 mg/kg administered i.p.). In the acute post-operative period following the contusion injury, all animals received extensive welfare checks and during this period animals were housed in soft-bedding and absorbent lined cages with less sawdust, to maximize ability to locomote, before returning to standard husbandry conditions.

#### Chondroitinase gene, lentiviral vector (LV-ChABC) and intraspinal injections

The *Proteus vulgaris* ChABC gene was modified with mutations targeted to remove five cryptic *N*-glycosylation sites and addition of a mammalian signal sequence and resynthesized with mammalian preferred codons to make a mammalian-compatible engineered ChABC gene (Muir et al., 2010). The modified ChABC cDNA was subcloned into a lentiviral transfer vector (termed LV-ChABC) with the mouse phosphoglycerate kinase (PGK) promoter, as previously described in detail (Bartus et al., 2014). The resulting vector was integrated, self-inactivated, and pseudotyped with VSV-G (vesicular stomatitis virus G). Viral particles were concentrated by ultracentrifugation and titred by a p24 antigen ELISA assay. The resulted viral titre used for both studies was 1.03E10 GC/ml (gift from Joost Verhaagen, the Netherlands Institute for Neuroscience). The animals designated to receive LV-ChABC treatment received two 0.5 μl intraspinal injections of LV-ChABC immediately after the injury, rostral and caudal to the contusion site (one injection per side). Injections were delivered at a rate of 200 nl/min using an ultra-micropump III (World Precisions Instrumentation, Europe). Following the intraspinal injection, wounds were sutured, and anaesthesia was reversed. All animals received post-operative analgesia (carprofen mg/kg) for three days, saline for rehydration (3–5 ml) for five days, and a 1 week course of antibiotics (baytril; mg/kg).

### Behavioural rehabilitation and assessment

The Whishaw window reaching and the Montoya staircase tasks were utilized for daily behavioural rehabilitation and to asses recovery of upper limb function. All rats were habituated and trained on the tasks for 4 weeks prior to any surgical procedures and baseline scores were obtained prior to electrode implantation and prior to contusion surgery (Fig. 1A). For the Whishaw window reaching tasks the animals were trained to repeatedly reach for sugar pellets (Test Diet, Sandown Scientific) through a slit in a Plexiglas cage. During the weekly behavioural assessment animals received a total of 20 pellets and were scored on how many they successfully retrieved and ate. For the Montoya staircase animals were placed in a chamber leading to a platform with access to a welled staircase on both sides where animals were trained to bilaterally reach for the sugar pellets placed in the wells. During the weekly behavioural assessment animals were scored on the number of pellets out of a total of 28 (14 per side) pellets were successfully retrieved and eaten. Coloured pellets were used to identify the levels of success in this task. The behavioural rehabilitation protocol was initiated at 7 dpi and animals performed both tasks daily (5 days/week) for 15 minutes.

**Figure 1.**
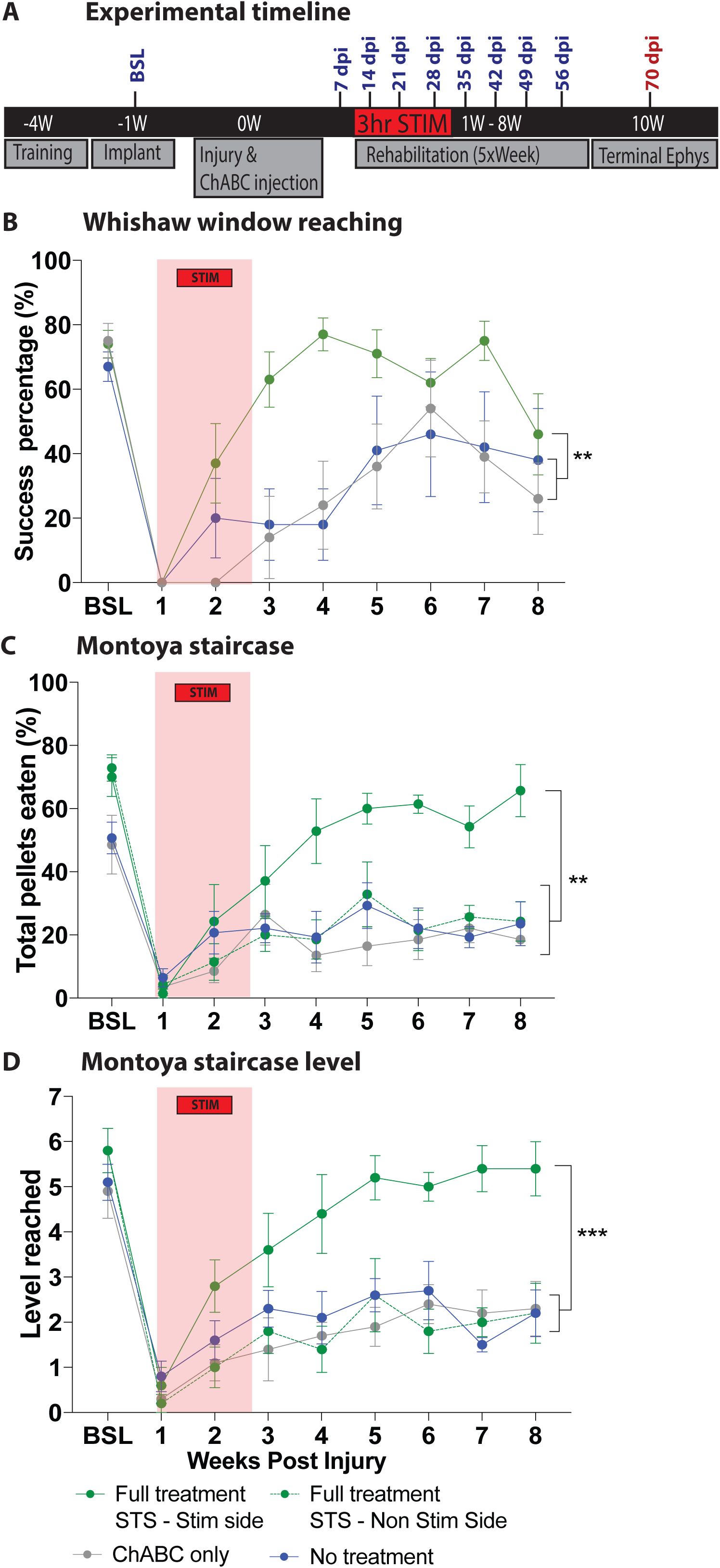
Experimental timeline and weekly behavioural task assessments. (A) Experimental study timeline (B) The Whishaw window reaching success rates indicates that the full treatment group (green) showed a significant improvement on regaining skilled reaching function (F16,64=2.379, p<0.0075 two-way RM ANOVA, Tukey’s *post-hoc*). (C) The Montoya staircase task measuring the number of total pellets eaten indicated that the animals in the full treatment group performed better with the side contralateral to stimulation (Stim side; green) compared to the side ipsilateral to stimulation (Non-stim side; green dotted line) (F24,120=2.304, p<0.0017 two-way RM ANOVA, Tukey’s *post-hoc*). Red coloured box indicates the time when the full treatment group received STS. (D) The Montoya staircase task measuring the difficulty level that animals from each group reached accurately for, demonstrating that the full treatment with STS stimulated side group accurately reached for the higher difficulty levels (F24,96 =2.634, p<0.0005 two-way RM ANOVA, Tukey’s *post-hoc*).

### Cortical epidural neuromodulation paradigm

The epidural cortical stimulation protocol was initiated 7 dpi and after all animals had fully recovered from the contusion injury surgeries. The animals were transferred from their home-cage to the stimulating chamber and acclimatized for 10 minutes before every session. The cortical stimulation protocol included M1 epidural stimulation for 3 hours for 12 consecutive days (5 pulses at 500Hz, 0.2 ms biphasic balanced pulses, every 2s) for all animals designated to a treatment group that included epidural cortical stimulation.

### Terminal electrophysiology experiments

Terminal experiments were performed following the last behavioural timepoint at 10 weeks post injury. Animals were anaesthetized (ketamine, 60 mg/kg, and medetomidine, 0.25 mg/kg) and the depth of anaesthesia was regularly assessed by monitoring pedal withdrawal reflexes and respiratory rate. Core temperature was maintained close to 37°C using a self-regulating heated blanket. During the experiment the animals were stimulated through the implanted cortical electrode at maximum current (10mA, 2 pulses, 0.2 ms biphasic pulses, every 30s) and muscle activity was recorded from percutaneously placed EMG electrodes on the bicep brachii muscle group contralaterally to the cortical stimulating electrode. The percutaneous EMGs were manufactured in house using a 27 gauge needle with a hooked wire on the tip and they were referenced to a ground wire inserted in the animal’s back muscles. To analyse the MEP recordings, EMG waveforms were first rectified and then filtered through a 5-point smoothing algorithm, a low pass filter (230 Hz, 310 decibels), DC offset correction and were then averaged. The area under the curve (AUC) of the motor evoked potentials (MEPs) was measured for each animal at maximum current (6-8mA) intensity.

### Perfusion and tissue processing

At the completion of experiments, animals were deeply anesthetized (sodium pentobarbital; 30mg/kg, i.v.) and perfused transcardially with heparin-containing saline, followed by 4% paraformaldehyde (in 0.1 M phosphate buffer; pH 7.4). For perfusion, a peristaltic pump was used at a pre-determined flow rate that depended on the animals’ weight (at 40-50%). The brain and spinal cord were removed, post-fixed in the same solution overnight and then transferred to PBS-azide for long term storage or 30% sucrose in 0.1 M phosphate buffer for cryo-protection respectively. Immediately after perfusion, lesion site tissue from the cervical C5-C6 vertebrae was dissected (10 mm with the lesion epicentre located centrally). Tissue was post-fixed for 2 h, cryo-protected in 20% sucrose for 48 h, then embedded and frozen in Optimum Cutting Temperature Compound before being cut into serial (20 μm) sections, cut transversely and tissue was mounted on microscope sections.

### Synaptic density immunohistochemistry and analysis

For immunohistochemistry, sections were blocked with either 10% normal donkey serum (Sigma Aldrich) in 0.2% Triton X-100 in PBS for one hour and then incubated overnight with a primary rabbit polyclonal anti-vGlut1 antibody (1:500; Synaptic Systems, 135 303). Then a species-specific secondary antibody raised in donkey (anti-rabbit Alexa Fluor 546 at 1:1000) was applied 24hr later. After three hours of incubation, the sections were thoroughly washed with PBS and coverslipped with Vectashield Antifade Mounting Medium with DAPI (Vector Laboratories Ltd). VGLUT1 stained sections were imaged at x10 and tile scans of the entire section were acquired. Imaging for the VGLUT1 synaptic density staining was done with a Zeiss LSM310 confocal microscope with consistent settings between all animals. Images were converted to 8-bit and rigid orientation-registered to a spinal cord template section corresponding to the appropriate spinal level by feature mark-up, and batch processed using custom macros for threshold analysis. A pixel value threshold was assigned as representative of positive immunostaining and maintained across all analysis for C4/5 region and again for C7/8 region. Pixels above this threshold were summed and averaged across two sections per animal for three animals per treatment group. Averaged threshold grey matter data per group were then converted to heat maps to display density of VGLUT1 positive staining using the NumPy and Matplotlib/Pyplot histogram2d functions in Python 3.6.

### Statistical Analysis

Statistical analyses were performed using Prism 7.0 software. For single time point recorded data statistical analysis was performed using one-way ANOVA with supplementary Tukey’s and/or Bonferroni’s post hoc post hoc tests, and for all chronically acquired data a two-way repeated measures ANOVA with supplementary Tukey’s and/or Bonferroni’s post hoc was performed. All data are presented as mean ± SEM values.

### Blinding and randomization

During surgical procedures the experimenter who performed the intraspinal injections was unaware of the animals’ identification numbers and to which treatment group each animal was assigned. All behavioural assessments were recorded on camera during the performance of the task and was then analysed by an experimenter blinded to treatment. The experimenter performing the electrophysiological terminal experiments was also blinded to each animal’s identification number and assigned group. Therefore, blinding was maintained as far as possible during data collection and analysis.

## RESULTS

### Short term stimulation combined with targeted rehabilitation and LV-ChABC leads to improved motor skilled function

We have previously shown that long term cortical stimulation (10 weeks) combined with daily rehabilitation and ChABC injections as part of a combinational treatment interventions leads to increased muscle function and recovery of skilled function after a cervical contusion injury (Sinopoulou et al. 2022). Here, we examined the effects of applying a short term (12 days) cortical stimulation paradigm as part of the same full treatment paradigm. All animals received 4 weeks of intensive training in skilled pellet reaching and then received a C5/C6 bilateral contusive injury (Fig. 1A). Contusion injury forces did not statistically differ between groups (Supp. Fig 1). Animals were assessed in the Whishaw window reaching task (Fig. 1B) and the Montoya staircase task (Fig. 1C and D). In the Whishaw window reaching task, the full treatment group showed an overall significant improvement compared to the single treatment (LV-ChABC only) and untreated groups (F16,64 =2.379, p<0.0075 two-way RM ANOVA, Tukey’s post-hoc) (Fig. 1B). Surprisingly, the group treated with the full combination of treatments showed an immediate improvement starting at 2 wpi which was increased until 5 wpi. However, at 6 wpi onwards the improvement effect started to diminish, and at 8 wpi the full treatment group showed similar reaching success levels to the single treatment (LV-ChABC only) and untreated groups. Furthermore, in the Montoya staircase task our results revealed that the stimulated forelimb (forelimb located contralaterally to the stimulated motor cortex) of the full treatment group showed a significant functional improvement in the task compared to the non-stimulated forelimb of the same group, the single treatment (LV-ChABC only) and untreated groups (F24,120 =2.304, p<0.0017 two-way RM ANOVA, Tukey’s post-hoc) (Fig. 1C). Interestingly, the stimulated forelimb of the full treatment group showed a functional improvement as early as 3 wpi compared to the non-stimulated forelimb, LV-ChABC only and untreated groups, which lasted until the chronic stages of the injury at 8 wpi, without any signs of the effect diminishing. The reaching difficultly level to which animals were able to accurately reach for was also examined in the Montoya staircase task (Fig. 1D). Similarly, to the amounts of pellets successfully reached, the stimulated forelimb of the full treatment group showed a significant improvement by accurately reaching for the lower wells, which translate to an increased difficulty level (F_24,96_=2.634, p<0.0005 two-way RM ANOVA, Tukey’s post-hoc).

### Terminal muscle activity recordings reveal an increase in evoked responses in the full treatment group

Terminal recordings acquired from the bicep brachii muscle group were obtained under anesthesia from intramuscular percutaneous recording electrodes placed contralateral to stimulation (as presented in the schematic shown in Fig. 2A). The area under the curve (AUC) of the evoked EMG responses was calculated and were then plotted for all the experimental groups, including a non-injured, not implanted control group. Our recordings revealed that the muscles on the stimulated forelimb of the full treatment group had a significantly larger AUC compared to the LV-ChABC only treated group, the untreated group and the non-stimulated side of the full treatment group. However, the full treatment group showed no statistically significant differences to the uninjured group, thus suggesting recovery of muscle activity (F_4,26_ =6.218, p<0.0012 one-way ANOVA, Tukey’s post-hoc) (Fig. 2B). Additionally, these results are also graphically represented in Fig. 2C.

**Figure 2.**
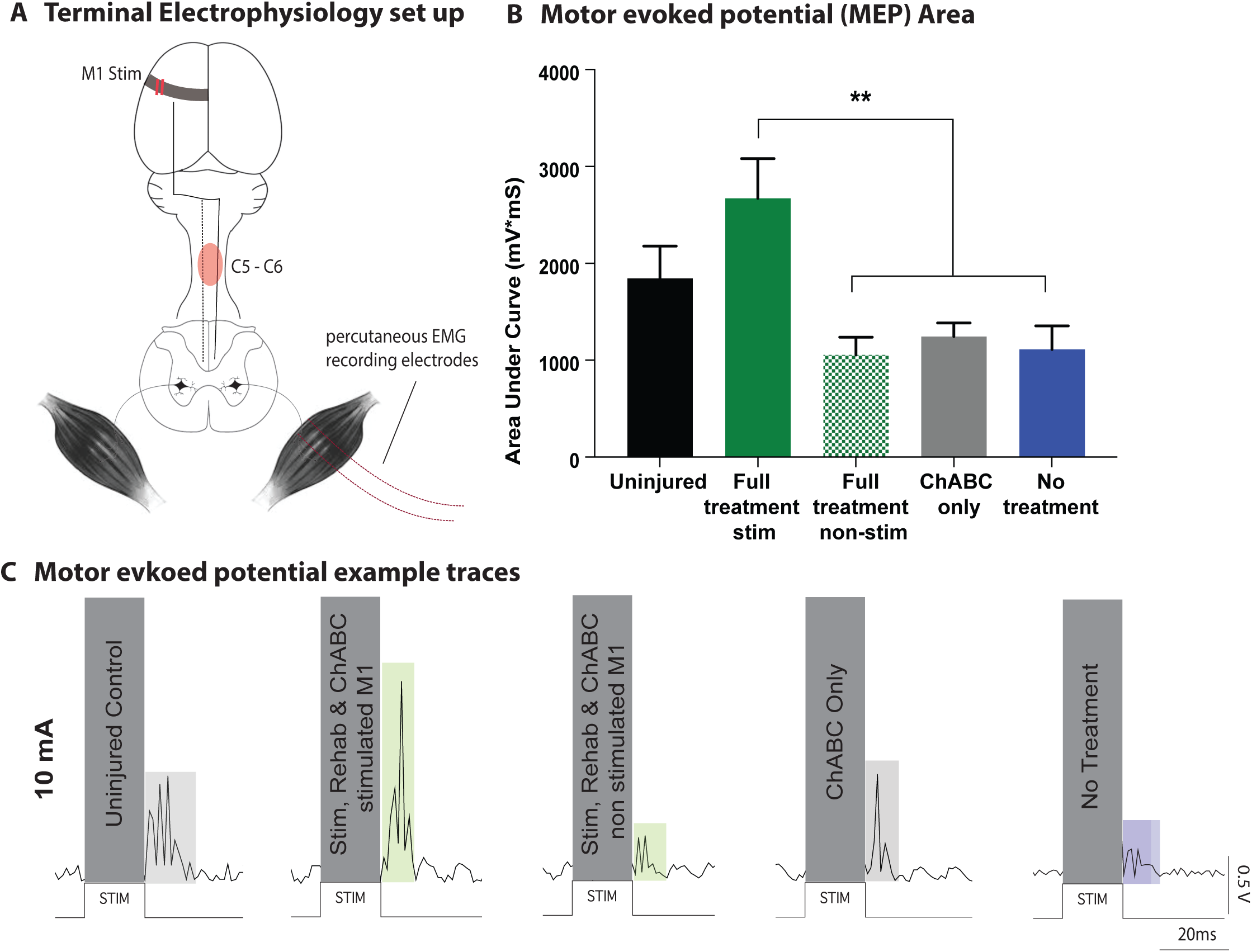
Terminal electrophysiology muscle recordings. (A) Schematic representing the terminal electrophysiology recording set up (B) MEPs recorded at 10mA under anaesthesia. The stimulated side of the full treatment with STS group had a significantly larger AUC (green solid bar) compare to all the other experimental groups except the uninjured group (F4,26=6.218, p<0.0012 one-way ANOVA, Tukey’s *post-hoc*). (C) representative traces from the bicep muscles obtained under anaesthesia from percutaneous EMG recording electrodes demonstrate similar results, with full treatment with STS group showing the largest evoked response.

### Overall synaptic density in the spinal cord revealed increased levels of VGLUT1 expression caudal to the injury level

To examine whether there was a change in the overall density of excitatory synapses after a cervical SCI and whether this was affected by the treatment paradigm administered, the expression of the presynaptic excitatory marker VGLUT1 was investigated in spinal cord sections rostral and caudal to the injury site. A threshold analysis was performed, representative heatmaps were produced and the density of VGLUT1 was quantified (Fig. 3). As it can be seen from the representative heat maps, VGLUT1 was expressed throughout the grey matter, particularly dense in laminae III-V (Fig. 3B and D). The pattern of VGLUT1 expression was similar for all treatment groups and the untreated group. Quantification revealed that rostral to the contusion injury at cervical levels C3/C4, the overall distribution of VGLUT1 was not significantly different between the experimental groups (F_2,15_ = 0.8792, p<0.4354 one-way ANOVA, Tukey’s post-hoc) (Fig. 3A). However, a significant difference in VGLUT1 expression within the experimental groups was observed caudal to the injury site at cervical levels C7/C8 (Fig. 3C). The full treatment group showed a significantly increased amount of VGLUT1 expression when compared to the untreated and the LV-ChABC only treated groups (F2,15 =3.927, p<0.0425 one-way ANOVA, Tukey’s post-hoc) (Fig. 3C). Heatmap representations in the full treatment group robustly show increased VGLUT1 expression in laminae III-V (Fig. 3D). The results suggest that the administration of different treatments after a contusion injury although it has no significant effect in the overall density of excitatory synapses rostral to the injury, caudal to injury is increased when the full combination of treatments is administered. Thus, our results reveal that the full combination of treatments has an effect on the overall density of excitatory synapses caudal to injury

**Figure 3.**
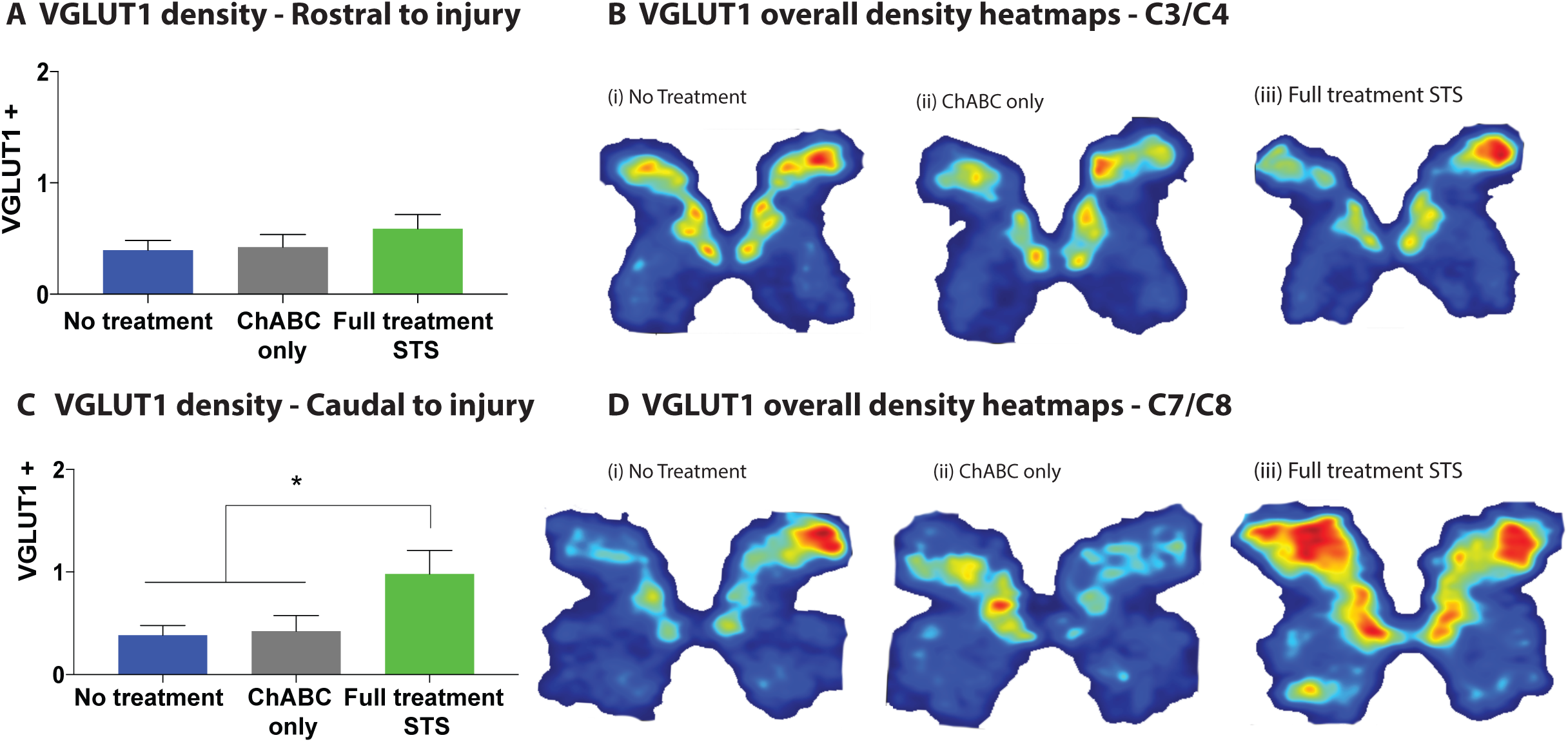
VGLUT1 expression rostral and caudal to injury. (A) Quantification of VGLUT1 positive pixels after the threshold analysis revealed no differences in the overall synaptic density of the spinal cord rostral to injury (F2,15= 0.8792, p<0.4354 one-way ANOVA, Tukey’s *post-hoc*). (B) VGLUT1 positive heatmaps representing a threshold analysis performed on the different experimental groups. (C) Quantification of VGLUT1 positive pixels after the threshold analysis revealed significant differences in the overall synaptic density of the spinal cord rostral to injury (F2,15=3.927, p<0.0425 one-way ANOVA, Tukey’s *post-hoc*) reporting that the full treatment group had significantly higher overall synaptic density compared to untreated group and the LV-ChABC treated group. (D) VGLUT1 positive heatmaps representing a threshold analysis performed on the different experimental groups.

## DISCUSSION

We showed that a novel neurorehabilitation paradigm consisting of short term cortical stimulation, upper limb focused rehabilitation and LV-ChABC intraspinal injections lead to significant recovery of function during skilled tasks such as the Wishaw window reaching and the Montoya staircase. We also report an increase in muscle activity equal to that of an uninjured animal when responses are evoked using epidural cortical stimulation under anaesthesia. Furthermore, we observed a significant increase in the overall amount of excitatory VGLUT1 positive synapses caudal to the injury in the spinal cord.

Physical rehabilitation has become an essential aspect of managing the effects of SCI and promoting recovery in humans, therefore it will be integral for any future therapeutic interventions to be considered in the context of physical rehabilitation. It was discussed previously that administration of LV-ChABC leads to a growth permissive environment and increases neuroplasticity in the spinal cord after a contusion injury (Bartus et al., 2014; James et al., 2015). However, it has been demonstrated that anatomical neuroplasticity does not always promote functional recovery (Wang et al., 2015). Therefore, it is essential to increase the functionality and connectivity of the newly formed connections by combing plasticity inducing agents, such as ChABC, with physical rehabilitation. Thus, it becomes feasible to harness the newly activity dependant sprouting. This has become a priority in the SCI preclinical research field (Zhao and Fawcett, 2013). Specifically, studies on C4 dorsal SCI rodent model (Wang et al., 2011) and a thoracic clip compression rodent injury (Alluin et al., 2014) demonstrated that animals who received ChABC treatment combined with task specific skilled reaching showed an increase in sprouting of the injured CST and with significant functional motor improvements, respectively. Similarly, past studies from our lab have reported a significant increase in skilled function post contusion injury after immediate LV-ChABC administration (Bartus et al., 2014; James et al., 2015) or dox-activated ChABC only (Burnside et al., 2018), in the present study we used LV-ChABC as part of a combination therapy, therefore a lower viral titer and a lesser amount of LV-ChABC was utilized (0.5 μl instead of 1 μl). Thus expectedly, the group treated with LV-ChABC only did not show a similarly robust effect as in our previous studies. Therefore, based on our results, we hypothesize that a lower titer although not effective to single handily promote functional changes, it was appropriate to create the necessary permissive environment for the stimulation and activity dependent changes to occur.

Furthermore, electrical neuromodulation has the capability to harness activity dependent structural plasticity and can potentially reactivate a growth program thus leading to regeneration after injury (Zareen et al., 2018). Here, we aimed to utilize this activity dependent regeneration and augment the effects using electrical neuromodulation in a permissive environment that has been created by administering LV-ChABC. Moreover, we showed an increase in excitatory synapses caudal to injury after administering the full combination of treatments. Our results support further evidence that has been previously reported on electrical stimulation suggesting that it can lead to an increase on the formation of new synapses by modulating the expression of Stat3 (Zareen et al., 2018), a pathway heavily involved in synaptic formation (Michalakis et al., 2013; Moon et al., 2013). Similar increase on VGLUT1 related synaptic density was also observed in a previous study from our lab that used a regulated ChABC delivery method and was positively correlated to functional recovery (Burnside et al., 2018).

Furthermore, other studies have suggested that rehabilitation can lead to activity dependent sprouting (ref). Here, our results revealed that the full combination of treatments lead to significantly restored skilled function in hand specific tasks. Specifically, in the Montoya staircase the full treatment group performed significantly better than the other groups and was also able to reach further in the more difficult level suggesting functional recovery. However, for the skilled reaching tasks, our results suggest that although short term stimulation can initially result in a significant functional improvement, this effect is not chronically sustainable for this type of specific task and it would potential require “boosting sessions”. In a clinical context our results reveal the potential combining of electrical stimulation of the brain, which is already being used along with rehabilitation after SCI, with other treatment interventions such as a neuroplasticity promoting agent.

## Acknowledgements

This work was supported by the International Spinal Research Trust (NRB113 to EJB and SBM) and the United Kingdom Medical Research Council (Senior Non-Clinical Fellowship Award G1002055 to EJB). We thank Naomi Roopnarine for technical support.

